# Exploring Rubiaceae fungal endophytes across contrasting tropical forests, tree tissues, and developmental stages

**DOI:** 10.1101/2024.02.13.580172

**Authors:** Humberto Castillo-González, Jason C. Slot, Stephanie Yarwood, Priscila Chaverri

**Author notes:** **Corresponding author**: Priscila Chaverri. Université Marie et Louis Pasteur, CNRS, Chrono-environnement (UMR 6249), F-25200 Montbéliard, France.

## Abstract

Fungal endophytes play a pivotal role in tropical forest dynamics, influencing plant fitness through growth stimulation, disease suppression, stress tolerance, and nutrient mobilization. This study investigates the effects of region, leaf developmental stage, and tissue type on endophyte communities in tropical plants. Young and mature leaves were collected from 47 Rubiaceae species, and sapwood from 23 species, in old-growth forests of Golfito and Guanacaste, Costa Rica. Fungal diversity and composition were assessed through metabarcoding of the ITS2 nrDNA region. Most identified ASVs belonged to the phylum Ascomycota. The orders Botryosphaeriales and Glomerellales significantly contributed to endophytic assemblages, without detection of host-specific communities. We observed significant differences in species richness across regions, confirming distinct compositions through beta diversity. No statistically significant variances were found between mature and juvenile leaf tissues. In contrast, leaves exhibited richer and more diverse assemblages than sapwood. As plants experienced diverse environments over time and space, our results may be influenced by changing structural and chemical properties through ontogeny. Given the potential impact of these fungi on agricultural and forest ecosystems, ongoing research is crucial to discern the roles of hosts, endophytes, and other ecological mechanisms in apparent colonization patterns.

## INTRODUCTION

Neotropical forests, which support a high diversity of terrestrial plants (1), host an extraordinary mosaic of microbial assemblages within plant tissues, driven by their provision of labile nutrients and niche space (2–5). These microbes play crucial roles in influencing host plants and driving forest dynamics, engaging in interactions that range from mutualism and commensalism to pathogenicity (6,7). Among these microbial groups, endophytes—microorganisms that reside in internal plant tissues without causing disease—are particularly noteworthy. Fungal endophytes offer numerous benefits to their plant hosts, such as enhanced immunity, improved growth, resistance to abiotic stressors, and the production of bioactive compounds (8,9). However, despite the significant diversity of fungal endophytes colonizing leaves and stems (4), they are traditionally understudied compared to root endophytes, mycorrhizae, and free-living forms. Consequently, much remains underexplored regarding their roles in natural ecosystems and their potential applications in improving crop health.

The mechanisms by which endophytic fungi colonize plant tissues remain a topic of research interest (1–3). Understanding the dynamics that determine their distributions and the environmental constraints shaping their assemblages is crucial. Evidence shows that broad-scale environmental factors such as temperature, elevation, precipitation (4,5), and forest type (6) significantly influence community composition. Additionally, genetic and geographic distance among host trees (7–9), as well as host growth stage and tissue type (10–12), also play crucial roles. Given these factors, endophytes that exhibit recurrence or specificity may play a significant ecological role in forests (13,14). Moreover, under the premise that high diversity maintains the overall integrity of an ecosystem (i.e., biological insurance hypothesis *sensu* Naeem & Li (15)) the hidden diversity of endophytes can enhance the fitness of individual trees and, by extension, shape forest composition and demography (16). Efforts to understand endophytic diversity are necessary for predicting their function in association with different tissues, hosts, specific environmental conditions, or entire ecosystems. In this sense, tropical forests offer an ideal setting for studying these patterns and interactions.

Rubiaceae is one of the largest and most complex families of angiosperms, comprising over 13,000 species that range from weedy herbs to large trees and occupy many habitats and biogeographical regions across all continents (17). However, they are predominantly found in the neotropics (18,19). In Costa Rica, historical data (20), listed 89 genera and 458 species present, but recent updates suggest that this count is underestimated. For example, the genus *Palicourea* alone, previously reported with 44 species, is now known to include 91 (21,22). This increased knowledge highlights the family’s extensive diversity. Noteworthy within Rubiaceae are several genera of cultural and economic importance such as *Cinchona*, *Coffea*, *Genipa,* and *Psychotria*.

Characterizing the fungal endophyte communities within Rubiaceae and the factors shaping their assemblages could provide unique insights into understanding their niche preferences in tropical forests. In this study, we investigated diversity patterns across a broad range of plant species, expecting to observe a vast taxonomic and functional diversity (7). We hypothesized that fungal endophytic communities would differ: (i) between forest regions due to varying environmental conditions; (ii) between individuals but showing similarities within closely related plants; and (iii) within a single tree, with differences between young and mature leaves, as well as between leaves and sapwood, exhibiting tissue-specific distributions. To further refine our hypotheses, we aimed to understand which fungal groups drove these differences by investigating whether certain endophytes exhibit cosmopolitan behavior or have significant associations with specific environmental conditions or host characteristics.

## METHODS

### Study site

Old-growth forests from two regions in Costa Rica were sampled. Specific sampling locations classified by climatic subregions are shown in Figure 1. Golfito, in the South Pacific, was visited in April 2017. All of the sampling sites in this area are characterized by typical tropical rainforest conditions, with no defined dry season, intense rainfall, and dense evergreen vegetation spanning two to three layers (23). In May 2017, sampling was conducted in Guanacaste, specifically in two distinct regions (North and North Pacific), comprising three subclimates: seasonal dry forest, semi-deciduous forest influenced by monsoons, and the forested slopes of a volcano, influenced by surrounding mountain systems (23). Climate data, including mean annual temperature, mean annual precipitation and elevation were obtained from the WorldClim database (Table 1) (24).

**Figure 1.**
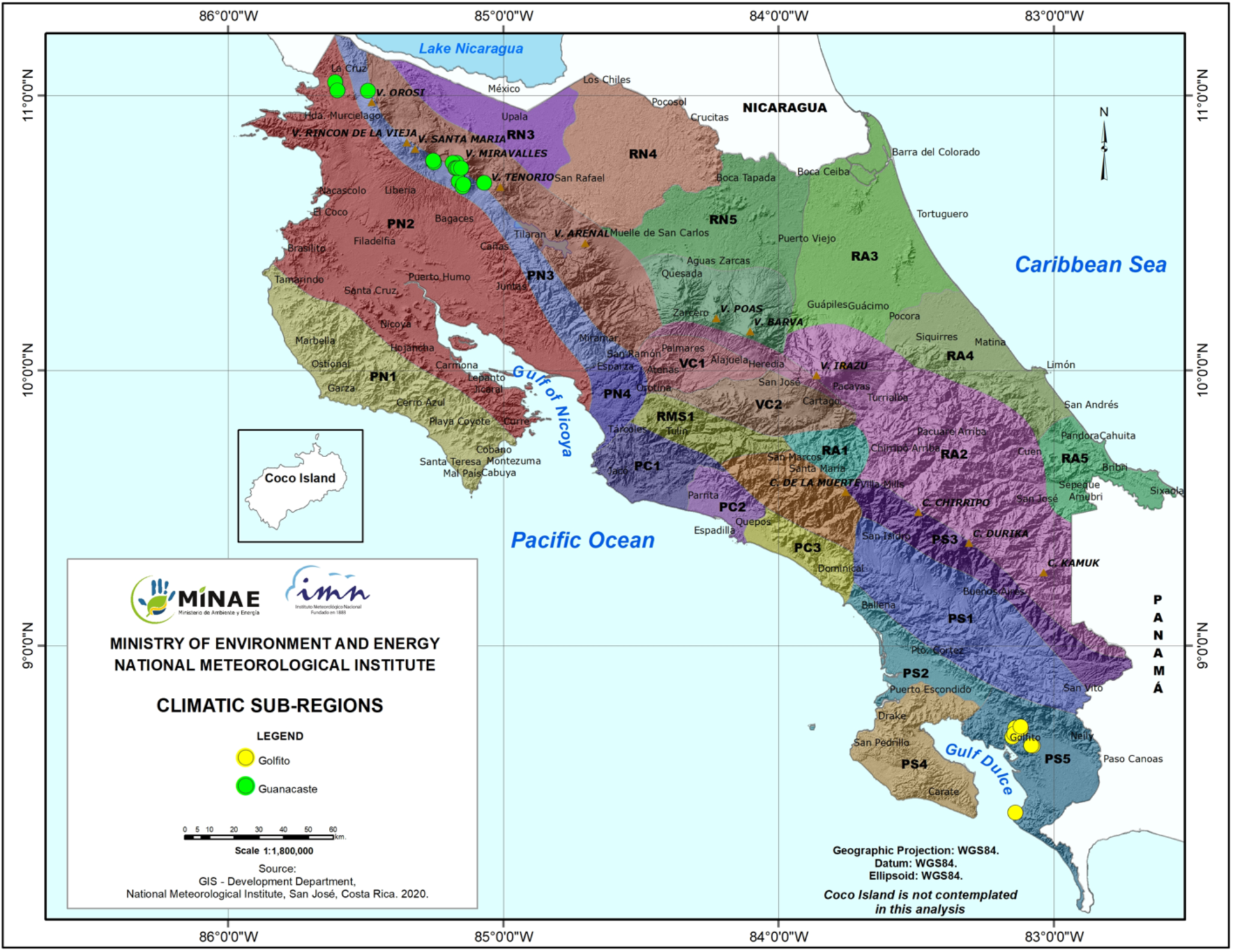
Map of Costa Rica showing climatic subregions and sample locations. Legend: RN1 – Eastern slopes of the Guanacaste and Tilarán mountain range (very humid subtropical forest), PN2 – North Pacific Central subregion (tropical dry forest), PN3 – Slopes and foothills of the Guanacaste and Tilarán mountain range subregion (monsoon-influenced rainforest), PS5 – Pacific slopes of the Talamanca mountain range (low montane rainforest). GPS locations for each site: **Guanacaste:** 10.668483 N, - 85.147333 W; 10.677167 N, - 85.143889 W; 10.681444 N, - 85.068889 W; 10.686972 N, - 85.161389 W; 10.735542 N, - 85.151997 W; 10.735675 N, - 85.163583 W; 10.735928 N, - 85.168861 W; 10.736069 N, - 85.173694 W; 10.754444 N, - 85.169244 W; 10.754583 N, - 85.182722 W; 10.755889 N, - 85.251833 W; 10.762333 N, - 85.254639 W; 11.01725 N, - 85.490778 W; 11.018633 N, - 85.601944 W; **Golfito**: 8.392333 N, - 83.13863 W; 8.635881 N, - 83.075394 W; 8.6367222 N, - 83.0815556 W; 8.668781 N, - 83.149222 W; 8.679278 N, - 83.143139 W; 8.698944 N, - 83.118911 W; 8.701139 N, - 83.138722 W; 8.706139 N, - 83.119631 W. Figure created by the National Meteorological Institute of Costa Rica at our request based on Solano & Villalobos (2001).

**Table 1.** Characterization of collecting sites, including mean environmental variables in locations (± SE).

| Location | Sites | Plants<br>(n) | Annual Mean<br>Temperature<br>(°C) | Annual<br>Precipitation<br>(mm) | Elevation<br>(m) |
| --- | --- | --- | --- | --- | --- |
| Guanacaste | 14 | 23 | $22.94 \pm 0.12$ | $2718.32 \pm 45.03$ | $749.56 \pm 41.72$ |
| Golfito | 8 | 27 | $24.65 \pm 0.19$ | $3522.09 \pm 21.17$ | $124.18 \pm 12.23$ |

### Sampling

Sampling of leaves and sapwood was conducted following a modified version of the protocol described by Gazis & Chaverri (25). Young and mature healthy leaves from 50 Rubiaceae individuals (distributed in 47 species from 27 genera, 10 tribes and 3 subfamilies) were collected (Supplementary Table S1). The taxonomy was assessed by a local botanist, and vouchers of all individuals were deposited in the University of Costa Rica Herbarium (USJ). We sampled plants as we encountered them, aiming to include as many different species as possible. For each individual sampled, fully expanded leaves were collected from three randomly selected branches, chosen from various parts of the crown. To differentiate developmental stages, we selected one newly emerged and one mature leaf per branch, totaling six leaves per plant.

For 23 of the individuals, which were woody trees with a minimum trunk diameter-at-breast-height of approximately 10 cm, a sterilized knife was used to cut a sliver of outer bark at shoulder height to expose the alburnum and one piece of living sapwood tissue ca. 30 × 5 mm was excised from the exposed area. All collected samples were put in a clean plastic bag and placed on ice for transport to the field workstation for immediate processing. A leaf section of ca. 20 × 4 mm was cut and surface-sterilized using sequential immersions in a 2% sodium hypochlorite solution (1 min), 70% ethanol (30 sec), and sterile distilled water (30 sec). Finally, leaves were dried with autoclaved tissue paper and stored in microtubes containing silica gel before being transferred on ice to the laboratory for DNA extraction (26). Sapwood samples were briefly singed with a flame to eliminate dust and potential surface contaminants introduced during handling, and then placed in microtubes.

### Sequencing

Samples were transferred to a pre-filled 500-micron Garnet and a 6-mm Zirconium Grinding Satellite bead tube (OPS Diagnostics LLC, New Jersey, USA) and ground using a FastPrep® bead mill (MP Biomedicals, Santa Ana, California, USA). Each tube was treated to a grinding cycle of speed: 6.0 m/s for 45 seconds, or until no visually recognizable fragments remained. Each sample (tissue section) represented a biological replicate, and no technical replicates were performed. Total DNA was extracted using the commercial kit Qiagen DNeasy Plant Mini Kit® according to the manufacturer’s instructions (Qiagen, Hilden, Germany). Genomic DNA was sent to Naturalis Biodiversity Centre (NBC, Leiden, The Netherlands) for targeted amplicon metagenomics (metabarcoding) of the ITS2 nrDNA region using Ion PGM Torrent^TM^ technology (Thermo Fisher Scientific, Waltham, MA, USA). The primers used were fITS7 and ITS4, which are fungal specific and with less amplification of plant ITS (27). To ensure accuracy in downstream analyses, a positive control with a non-Costa Rican *Russula galochroa* DNA and a negative control (blank) were included throughout. According to NBC, PCR was performed in a BioRad C1000 thermocycler (Hercules, CA, USA), quality of the PCR products was checked with a QIAxcel (Qiagen), PCR products were cleaned with NucleoMag NGS Clean-up and Size Select magnetic beads (Takara Bio Inc., Japan), the pool was run in an Agilent Bioanalyzer (Santa Clara, CA, USA) using a DNA High Sensitivity chip and amplified on the Ion One Touch system (Ion PGM^TM^ Hi-Q^TM^ View OT2 kit), the results were measured on the Qubit (ThermoFisher) for the Ion Sphere^TM^ Quality Control kit (ThermoFisher), and finally the sequence run was done on the Ion PGM Torrent using the Ion PGM^TM^ Hi-Q^TM^ View Sequencing kit. The resulting raw data (fastq files) are deposited in GenBank under the BioProject identifier PRJNA889378.

### Fungal community analyses

#### ASV classification and taxa assignment

Primers and barcodes were trimmed from each read using the command line tool ‘cutadapt’ (28). Subsequent processing was performed in RStudio v. 2022.12.0+353 (29) with R v. 4.0.3, employing the DADA2 v1.18.0 pipeline (30). Raw reads were subjected to data curation aimed at identifying and removing spurious contigs through various filtering processes. For instance, chimeric sequences were removed. Reads with total counts below 0.001% of the total reads and singletons were excluded. Quality filtering included a maximum expected error rate of 2 for both forward and reverse reads, and sequences were trimmed to a minimum length of 100 bp (36). Amplicon Sequence Variants (ASVs) were assigned taxonomically by comparing them to the UNITE database (36) v8.3 (31) training set from the DECIPHER package, v 2.18.1 (32). Only sequences classified as fungi were retained. Additional independent filtering steps included the exclusion of samples with total read counts below 0.001% of the total reads, and taxa with low variance across samples (variance threshold of 1e-5). A prevalence threshold of 1% of total samples was applied, retaining taxa appearing in at least this proportion of samples. The accuracy of the pipeline was validated using the positive controls, and sequences from negative controls were removed to ensure dataset integrity.

#### Diversity and community structure analyses

Statistical analyses and plotting were conducted using the phyloseq v1.38.0 (33), vegan v2.5.7 (34), microbiome v1.12.0 (35) and ggplot2 v3.4.1 (36) packages. Endophyte alpha diversity for each dataset was estimated using the Shannon, Simpson, and Chao1 indices for each variable and then transformed to Hill Effective Species Numbers (37,38) for comparisons by Mann-Whitney U test. This transformation integrates species richness and abundance distribution, adjusts sensitivity to rare versus common species, and provides a consistent measure of diversity (43,44). It also allows for more robust comparisons across sites with varying numbers of species and levels of evenness. To visualize community patterns (beta diversity), non-metric multidimensional scaling (NMDS) was performed using the ‘metaMDS’ function from the vegan package with the ‘bray’ method and 100 random starts. For statistical analyses, data were normalized by total count per sample, converting them into proportions (relative abundances), and Bray-Curtis distances were calculated with the ‘vegdist’ function. Homogeneity of dispersion was assessed with the ‘betadisper’ function, followed by a permutation test using ‘permutest’. Statistical differences in endophyte assemblages across variables (forest regions, tissue types, developmental stages) were evaluated with the ‘adonis2’ function for PERMANOVA.

Relative abundance at the family level was calculated. We retained groups contributing to more than 1% of the average relative abundance across the six categories: forest regions (Golfito and Guanacaste), tissue types (leaf and sapwood) and developmental stages (young and mature).

These groups are referred to as the ‘top families’ in downstream analyses. The overall prevalence (i.e., the number of samples in which the taxa appear) of these families was computed. To estimate how strongly they deviate from a random preference among the plant partners available at the study sites, we calculated the d’ index of specialization using the bipartite v 2.18 package (39), function ‘dfun’. ASVs were statistically classified as generalists or specialists based on their habitat affinity using a multinomial species classification method (CLAMtest) in vegan. Indicator taxa (ASVs with p-values < 0.05) from the six categories were identified using the ‘multipatt’ function in the indicspecies v1.7.12 package (40). Methods regarding Species Accumulation Curves, Mantel Correlation Tests, Distance Decay, Canonical Correlation Analysis and the assignment of functional traits are described in Supplementary Methods S2.

## RESULTS

A total of 6,467,267 reads passed the quality control filters. The mean number of sequences per sample was 58,465 for leaves and 26,989 for sapwood. We identified 3,218 putative taxa (ASVs) belonging to at least 232 different genera (Supplementary Table S3). ASV identification rate was 67%, 50% and 33% at the order, family, and genus levels, respectively. The species richness or number of unique ASVs recovered per sample (triplicates, in the case of leaf tissue) ranged from 2 to 337, with a mean of 67.

All datasets had a high number of single occurrences, and no asymptote was reached in the species accumulation curves (Supplementary Figure S4) indicating that the datasets are incomplete, and the number of samples was not sufficient in capturing the expected endophyte diversity (Hill’s q = 0). However, when focusing on dominant species only (Hill’s q = 2), all curves, except for sapwood, reached a plateau.

Fungal endophyte assemblages differed between forest regions and tissue types (Table 2). Samples originating from Golfito exhibited consistently higher alpha diversity indices compared to those from Guanacaste, as supported by a Hill numbers comparison (q=0,1,2) that demonstrated statistical differences for q=0 (p ≤ 0.05) and q=1 (p ≤ 0.01). Notably, leaf and sapwood tissues were found to harbor distinct fungal communities, with leaves exhibiting significantly richer assemblages (p ≤ 0.001 in all instances). Regarding the developmental stage, the taxa richness index (q=0) was significantly higher in mature leaves compared to new leaves. In contrast, the other two diversity indices (q=1, q=2) showed slightly greater diversity in new leaves, though these differences were not statistically significant.

**Table 2.** Alpha diversity represented by Hill Effective Species Numbers (± SE) for the evaluated variables. Hill numbers represent taxa richness (Chao1), Shannon diversity (the exponential of Shannon entropy), and Simpson diversity (the inverse of Simpson index). Mann-Whitney U test p-values reported.

| Variable | Category | Sample size | Obs. Species richness | Shared number of ASVs | Taxa richness (q = 0) | Shannon diversity (q =1) | Simpson diversity (q = 2) |
| --- | --- | --- | --- | --- | --- | --- | --- |
| Location | Golfito | 66 | 2146 | 315 | 79.52 ± 7.91 | 25.41 ± 2.26 | 13.75 ± 1.08 |
|  | Guanacaste | 57 | 1387 |  | 51.81 ± 6.69 | 20.06 ± 2.38 | 11.48 ± 1.21 |
|  | p-value |  |  |  | **0.00655** | **0.04821** | 0.07629 |
| Tissue type | Leaf | 46 | 1514 | 96 | 61.89 ± 7.03 | 21.22 ± 2.61 | 11.84 ± 1.38 |
|  | Sapwood | 23 | 240 |  | 11.83 ± 1.64 | 1.57 ± 0.17 | 0.66 ± 0.05 |
|  | p-value |  |  |  | **4.99E-06** | **8.22E-06** | **2.41E-04** |
| Development stage | Mature | 50 | 2253 | 912 | 92.40 ± 8.56 | 22.80 ± 2.51 | 14.31 ± 1.22 |
|  | New | 50 | 1756 |  | 66.18 ± 7.87 | 25.62 ± 2.63 | 14.56 ± 1.31 |
|  | p-value |  |  |  | **0.0067** | 0.4628 | 1.0000 |

Using non-metric multidimensional scaling (NMDS) we observed differences between location and dispersion of points, with Guanacaste samples appearing more scattered (Figure 2A); three outliers were removed for visualization purposes. Community dissimilarities between these variables were corroborated via PERMANOVA, which indicated significant separation in fungal assemblages when comparing between the different forest regions (p < 0.001). In the case of tissue type, points from leaf and bark communities were dispersed differently in the NMDS analysis (Figure 2B) and presented significant heteroscedasticity (p < 0.01), which has been shown to affect PERMANOVA testing (p < 0.001). No clear location effect was observed in the NMDS plot; however, the dispersion of the points varied: leaf samples converged on the interior of the cluster while sapwood samples, being more variable, were found mostly around it; two samples were removed for better visualization (figure 2B). Considering just the leaves at the two different development stages, no effect was detected, there was a high overlap between the fungal communities (Figure 2C), and the PERMANOVA indicated no significant differences (p = 0.8122). Fungal endophyte communities did not cluster based on individual plants or at any higher taxonomic level (i.e., genus, tribe, subfamily). The pattern for tribe is shown in Figure 2C, other levels of taxonomy are depicted in Supplementary Figure S5.

**Figure 2.**
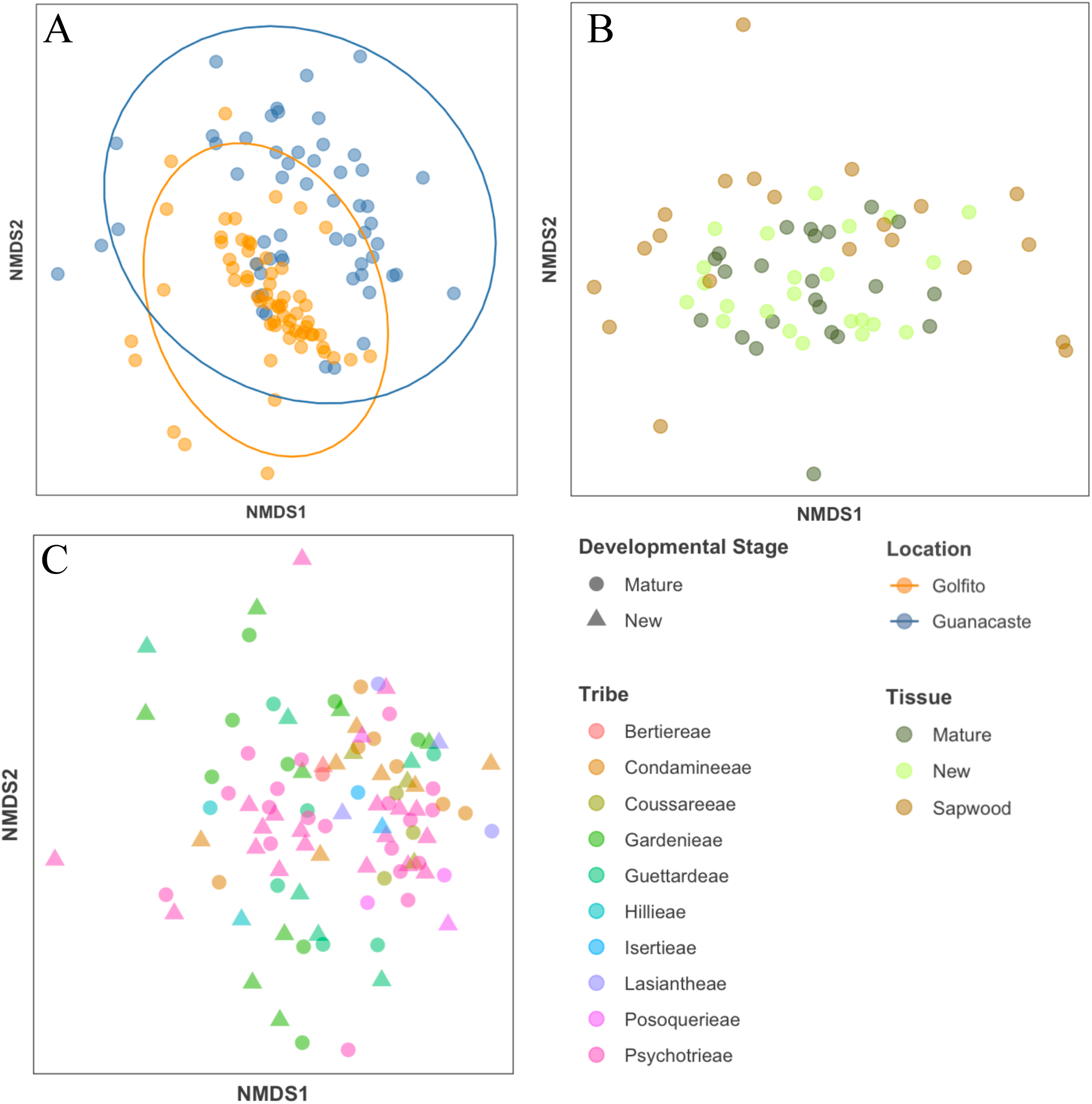
NMDS of Bray Curtis distances showing dissimilarities between the structure of endophyte community assemblages from (A) different locations, (B) different tissues and (C) different plant tribes, developmental stage represented here by shape.

The phylum Ascomycota was the most abundant, representing 83% of all fungal ASVs. Within this group, the most abundant and prevalent classes were Sordariomycetes followed by Dothideomycetes. Tremellomycetes and Agaricomycetes were the most abundant and prevalent Basidiomycota, making up 8.2% of the ASVs. We characterized endophytic assemblages dominated by a few abundant dominant taxa and many rare taxonomic groups. For example, we identified 156 different families (Supplementary Table S6) from which Glomerellaceae, Phyllostictaceae and Mycosphaerellaceae composed the bulk of biodiversity (45.14%) present within the endophytic assemblages (Figure 3).

**Figure 3.**
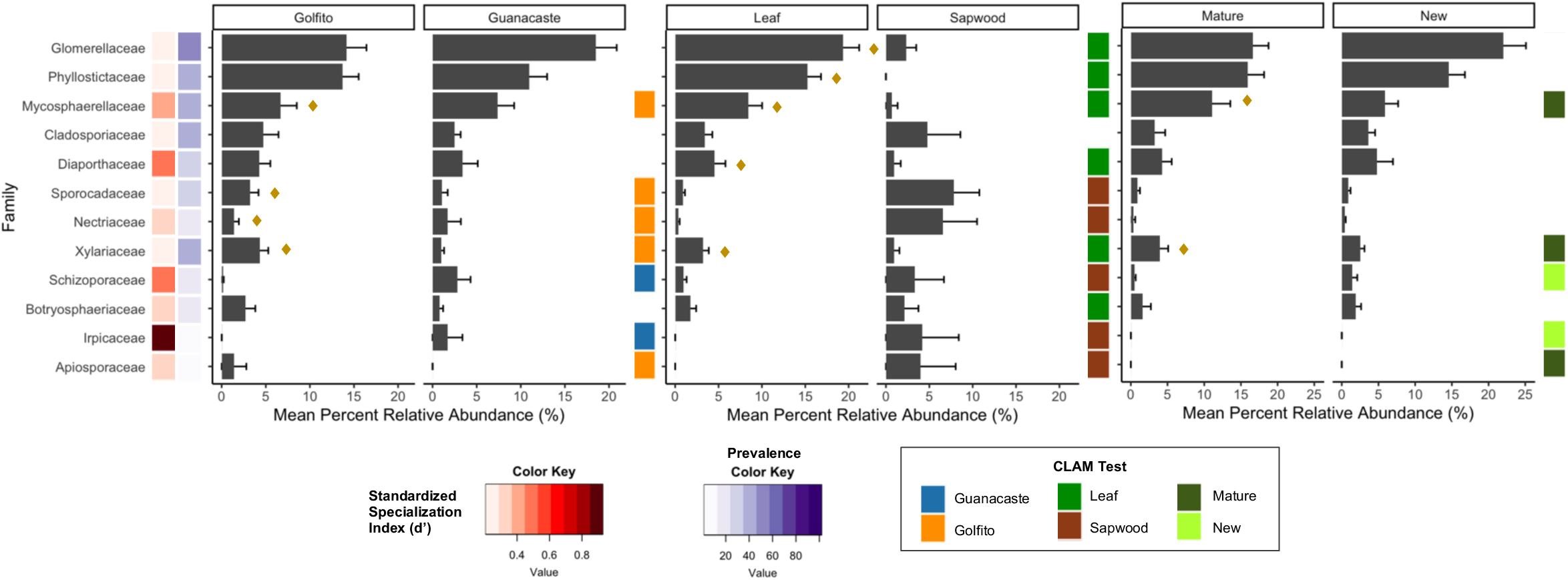
Prevalence and mean percent relative abundance of the 12 most abundant families weighting all habitats within location, tissue type and foliar developmental stage. Families are organized based on abundance, high to low, top to bottom. Mean abundance is plotted with error bars representing the standard error. d’ index calculates how specialized the identified taxa are in relation to the 47 plant putative species. Denoted with a diamond symbol (♦) are the families with indicator species within the different habitats; and habitat specialization is indicated by the CLAM test.

The most abundant fungal families were not the most prevalent in all cases (e.g., Xylariaceae), but they tended to correspond (Figure 3). The top 12 families comprised 59% of the total relative abundance. With the d’ index, we determined that the most prevalent and abundant families (i.e., Glomerellaceae and Phyllostictaceae) had values closer to 0 (no specialization). This indicates that members of these fungal families interact with a broad host range. In contrast, Irpicaceae, had a low prevalence and presented a value closer to 1 (perfect specialist). Interesting is the case of Apiosporaceae, that despite a low prevalence, showed little specialization. The major fungal families were similar between Golfito and Guanacaste (Figure 3), but some of the groups that presented very low abundance in Golfito, such as Schizoporaceae and Irpicaceae, were more abundant in Guanacaste and such preference was highlighted by the CLAM test (Figure 3).

We did not find indicator species among the top 12 families in the sapwood communities. In contrast, some indicator species were identified for leaves. For example, Phyllostictaceae, despite being the second most abundant and prevalent family overall, was not found in sapwood. Sporocadaceae and Nectriaceae, both of which had very low abundance in the foliar tissues, comprised around 10% of the relative abundance in sapwood samples. As a matter of fact, most families with less than 5% relative abundance in leaves clearly presented higher values in sapwood, except for Diaporthaceae and Xylariaceae. The CLAM test classified most families as specialists in one or the other habitat, except for Cladosporiaceae, which happened to be the only group that did not show habitat preference in any of the evaluated instances (Figure 3). Interestingly, some of the families preferred sapwood over leaves but displayed a preference for a developmental stage when found in foliar tissues: Schizoporaceae, Irpicaceae, and Apiosporaceae. Within developmental stage, less than half of the families showed habitat preference or specialization and only two were recognized as indicators for mature leaves and none for new. Some of the families exhibited habitat preference in all the evaluated instances, for example, Mycosphaerellaceae and Xylariaceae preferred Golfito, leaf tissue, and mature stages; while Schizoporaceae and Irpicaceae preferred Guanacaste, sapwood, and new leaves (Figure 3).

## DISCUSSION

The diversity of fungi found in the leaves and sapwood of tropical plants belonging to the Rubiaceae family did not show discernible patterns of specific host affiliation. Highly abundant community members seemed to be consistently present within the microbiome across different habitats and diverse host species. Fungal communities in leaves appear to be stable in early tissue development and persist in mature stages. As expected, tissues contained distinct habitats for symbionts. The mechanisms of community assemblage at regional scales are not clear. We found weak evidence for dispersal limitation and environmental partitioning; therefore, the observed patterns may be a result of unknown environmental variables that are correlated with the different geographic locations.

### Cosmopolitan fungi dominate hyperdiverse endophyte communities

A fungus’ niche preference is largely determined by the interaction between genotype and environment (41). Therefore, it can vary among forests that differ in plant community composition and biogeographic and environmental conditions. Communities of endophytes with a broad host range may be completely shaped by the environment, while for those with host restrictions the plant might exert a stronger influence (42). Results from this study support previous findings that in high-diversity natural systems, such as tropical forests, the endophyte communities are typically composed of numerous taxa of low relative abundances (43), and tend to be dominated by few fungal taxa with a broad host range. The latter are predicted to have a widespread availability of suitable partners, displaying loose host affiliation, and occupying geographically and taxonomically disparate plant hosts (44–46), patterns also found in this study (Figure 3).

Host jumps or shifts by microbial symbionts usually occur among closely related host taxa and are driven by the exploitation of new adaptive zones (47). Endophytes may display host preferences at the plant family (48) or order (49) level. In our study, we focused exclusively on samples from a single plant family, Rubiaceae, and found that taxonomy did not predict fungal community clustering (Figure 2C, Supplementary Figure S5). However, this does not exclude the possibility of host effects in other contexts, at a finer scale (e.g., within genus) or with more individuals per plant species (50,51).

### Forest regions shape fungal endophytic communities

The results of our study support the hypothesis that fungal endophytic communities differ between forest regions. Samples from the tropical rainforests of Golfito exhibited higher alpha diversity compared to those from Guanacaste (Table 2), which includes sampling sites classified as tropical dry forest. Notably, these ecosystems experience significant seasonal changes, including extended periods of drought. These shifting conditions lead to the presence of semi-deciduous trees, which alters the availability of host tissues and results in fluctuations in endophyte activity and diversity. For instance, some taxa may become dormant during dry periods, potentially leading to an underestimation of the local pool of endophytes. The temporal distribution of endophytes, documented in both temperate (52) and, more recently, tropical forests (44), supports this phenomenon. Differences between the regions were also evident in community similarity.

Moreover, our analysis revealed that endophytic communities in Guanacaste exhibited a more scattered pattern, suggesting higher intra-regional variability among those samples (Figure 2A). This observation aligns with and may be related to the variability in environmental conditions across the region (Table 1; Figure 1).

The environment can serve as a filter, regulating which particular taxa can colonize and persist within hosts (42,53). However, in this study, we did not find strong associations between environmental variables (i.e., elevation, temperature, or precipitation) and fungal community composition (Supplementary Figure S7, Supplementary Table S8). This suggests that environmental factors may not be the primary drivers of community variation. Instead, biotic processes such as priority effects may be more predictive of community structure. For instance, horizontally transmitted endophytes must compete with already established horizontally or vertically transmitted microbes, thereby restricting the pool of species able to colonize a given host.

Additionally, neutral processes, such as dispersal limitation and ecological drift, also play a significant role in the distribution pattern of endophytes (54–56). Successful host colonization and taxa distribution depend on factors such as dispersal opportunities and movement restrictions across different environments (57). Consequently, the similarity in species composition between ecological communities typically decreases with increasing distance, a phenomenon known as distance decay (58). Although dispersal limitation is not considered strong for endophytes, distance decay has been reported with varying magnitudes (59–61). Our study found a marginally significant interaction between geographic location and community structuring (Supplementary Table S8). Moreover, the endophytic communities were more strongly correlated with the geographic distance than environmental parameters (Supplementary Table S8), despite no clear evidence of distance decay (Supplementary Figure S9).

Both Golfito and Guanacaste shared the same dominant generalist taxa (Figure 3), characterized by high prevalence and abundance across a variety of plant hosts. In contrast, some fungi exhibited specialist behavior, such as Irpicaceae in Guanacaste samples, and Apiosporaceae in Golfito. This suggests that while generalists are broadly distributed, specialized interactions are more localized, likely due to habitat- and niche-specific factors. For instance, the availability of suitable plant hosts in a given ecosystem may limit the geographical distribution these specialized taxa (57,62). This differentiation underscores the role of niche processes in shaping fungal communities, leading to variations in both taxonomic composition and functional traits. At this broad spatial scale, environmental heterogeneity between regions may influence trait values more than to taxonomy, producing community variations that are not solely explained by taxonomic classifications (63,64).

### Tissue type predicts endophyte community variations

The phyllosphere is a dynamic and heterogeneous environment, whose surfaces are often considered relatively oligotrophic (65). The ability to colonize plant tissues is largely conditioned by complex physical-chemical interactions with their hosts. For example, fungi must pass through several lines of defense (66) and resist the host production of antimicrobial compounds (54,67). Results from the present study support our hypothesis, and previous studies, that endophyte communities are structured by plant tissue (43,68,69). We observed large differences in endophytes between leaves and sapwood. Leaves harbored more robust and rich assemblages than sapwood, although the latter is considered rich in organic nutrients and is a target of many different organisms (70). We observed little overlap between foliar and sapwood endophytes, suggesting spatial heterogeneity across plant compartments (71–73).

While most of the top families of fungi were present in both tissues, they differed in relative abundance and were not dominated by the same taxa. The most abundant family in sapwood was Sporocadaceae, and not surprisingly, both families belonging to the Basidiomycota in the top 12 families preferred sapwood according to the CLAM test (Figure 3). In the present study, Phyllostictaceae, the second most abundant family on the dataset was not detected in sapwood, indicating an affinity of this taxon towards foliar tissues. Meanwhile, other families (i.e., Nectriaceae, Apiosporaceae and Irpicaceae) almost imperceptible in leaves, presented higher abundance in sapwood, suggesting that tissue type correlates with selective enrichments of specific endophytes (74,75), and some fungi may not be able to persist or disperse between different tissues.

Specificity of fungal endophytes in bark has been previously reported (76). These fungi hold great but underappreciated ecological significance, for they inhabit large surfaces often targeted by insects and pathogens (77,78) and represent a passive reservoir of spores and latent life that will eventually play an important role in wood decay dynamics (79,80). Several studies on endophytic community profiles suggest broad correlations between diversity and select defense mechanisms in bark (81), e.g., reaction zones, the vascular cambium, and the transpiration stream in the sapwood (82). Furthermore, the presence of certain fluids, such as sap, can also act as a barrier depending on their levels of specific compounds (83,84). The sapwood of trees consists of dead xylem cells, and the presence of air-filled vessels and pits can create physical barriers that limit endophyte movement within the tissue (85,86). Similarly, the negative pressure within the xylem can hinder endophytes from colonizing this tissue (87), as they may not be able to penetrate the vessel walls or survive under conditions of low oxygen and nutrient availability and high-water stress (88,89). Therefore, bark acts as an environmental filter structuring inner bark fungal communities (79) which might explain the differences in community richness, diversity and composition observed in this study.

The canopy cover of trees creates dynamic microclimates that affect endophyte distribution and growth by modifying temperature, humidity, and wind conditions (90–92). Fungi release spores from within and upon canopies, that are then dispersed onto vegetation and exposed to various elements (93). During precipitation, for example, natural events such as throughfall and stemflow transport bioaerosols and spores through different habitats within a plant microbiome, including bark and leaves (94). The accumulation of endophyte propagules is then influenced by surface area, as plant structures of various sizes interact differently with rainwater and wind (95,96). Woody stems can be challenging to saturate with water due to their fiber content, cylindrical shape and thickness (97,98). Additionally, the large surface area on certain tree species, often covered in small grooves or ridges, increases the number of accumulation areas, such as fissures, cracks, tree hollows, and dendrotelmata (99,100). These structures enable the storage of a vast number of spores and serve as reservoirs for fungi, offering wide niches for their survival (101). In fact, a recent study found that large water-insoluble particles (e.g., conidia, spores) tend to accumulate more on bole surfaces than on leaves or branches (102). Although these spaces and other structures, such as lenticels, may act as entry points for fungal endophytes, complex polymers in this tissue, including lignin and suberin, pose significant barriers to endophyte colonization (103–105).

In foliar tissue, water droplets containing spores can adhere to trichomes, epidermal cell grooves, and the curvature of leaves (94), and use stomata, trichomes, and other structures to enter the tissue (106–108). However, the hydrophobic nature of the cuticle causes aerial depositions to be washed off during rainfalls, leading to self-cleaning surfaces (94). Despite leaves being rich in bioactive compounds with antifungal properties such as phenolics, alkaloids and terpenes (109,110), endophytes thrive in these environments due to high moisture content, and nutrient availability.

### Leaf developmental stage does not drive endophyte community differences

Generally, newly emerged leaves in woody plants start endophyte-free but rapidly become colonized by fungal spores (111). In this study, differences in foliar endophyte richness due to leaf developmental stage (young vs. mature) were not consistent across diversity indices (Table 2). We predicted that new leaves would present lower endophyte abundance due to hypothetical higher concentration of chemical defenses (112–114). Ontogeny-related shifts in endophyte diversity could also be anticipated due to various changes in tissue characteristics, including genetic traits, chemistry and topology (65), plant hormonal or physiological properties, and physical characteristics, such as tissue structure and geometry (115). However, it is important to note that these factors were not directly measured.

Our beta diversity analyses (Figures 2B, 2C) show that leaf developmental stage was not a strong predictor of the observed community compositions. Defense syndromes in plants are usually associated to strategies of resource conservation vs resource acquisition (e.g., young vs mature leaves) (116). In fact, some studies have found large qualitative and quantitative differences in metabolomic composition between expanding and mature leaves (117) (i.e., Optimal Defense Theory); however, this is not universally observed (118). Research in tropical trees has found low variation in defense traits and weak shifts in secondary chemistry during leaf maturation (110). This aligns with the “grow fast, die fast” hypothesis, which suggests that plants in these biomes adjust carbon allocation to balance early growth against defenses due to shorter leaf lifespans (119,120). Similarly, the resource availability hypothesis suggests that plants in resource-limited environments (e.g., boreal regions) invest more in constitutive defenses than those in resource-rich habitats (e.g., tropical regions) (121). This implies that plants in the tropics may not necessarily experience strong selective pressure to evolve higher levels of chemical defense against biotic threats (122).

Consequently, tropical plants may not exhibit significant variations in the phytochemical profiles of young versus mature leaves. This could account for the absence of significant differences in endophyte communities across developmental stages, as we observed no increase in endophyte relative abundance with tissue maturity. Additionally, both habitats were dominated by the same families, which can be considered prominent components of foliar endophyte communities in Rubiaceae tropical plants and are likely to play a significant role in ecosystem functioning. The CLAM test identified Xylariaceae and Mycosphaerellaceae as the two families that displayed a preference for the mature leaves’ habitat. These families are commonly known as foliar endophytes, but they also include pathogens, litter decomposers, and wood saprotrophs (Supplementary Figure S10).

Endophytes interact with both the internal environment of the host plant and the broader external abiotic conditions, that influence their distribution and functions in complex ways (123). We observed intriguing patterns in the composition and diversity of foliar endophytes in tropical Rubiaceae. However, important caveats must be considered when interpreting these findings. Although our analysis accounted for certain environmental parameters, we did not measure other significant factors such as shade and light availability (124,125), humidity (126,127), wind (128), ground topography (129), and soil characteristics (e.g., carbon content, texture, and physicochemical properties) (130,131). For instance, stable and functional soils, like those of tropical forests, are likely to host robust microbial reservoirs (132), facilitating the horizontal transfer of endophytes to the phyllosphere, adding another layer of complexity to plant-endophyte interactions.

Moreover, when accounting for geographic factors and habitat effects, it is crucial to acknowledge the potential influence of other unmeasured variables, including forest structure and composition, plant density (111), tree diversity, climatic gradients (123,133), and landscape-scale elements (134). Temporal factors, such as forest succession and seasonal variations also contribute (135–137). Moreover, the host’s age (138), genetics, architecture and spatial structure (111,139) introduce additional complexity that we did not capture. Nonetheless, the diverse nature of the observed endophytes implies that no single ecological effect is universal. While the determined patterns may be constrained by the specific environmental and climatic factors herein considered, endophyte-plant interactions represent a novel dimension of niche differentiation for coexisting tree species (132), still highly unexplored and nuanced by the intricate and context-dependent trade-offs shaping these communities.

With the imminent threat of climate change and the constant challenges in ecosystem management and agriculture, endophytes have the potential to transform the way we think about and manage plant stress resistance (140–142). Understanding their distributions and functioning in association with different host plants and ecosystems, and their response to environmental disturbances, is key to predicting the potential impact of many of these fungi in managed and natural ecosystems. How to translate this acquired knowledge into practical applications for forest conservation and agricultural practices remains a challenge. Many cryptic and undiscovered taxa, whose ecological roles are still unknown, await exploration. In the future, functional group and single trait approaches could offer valuable insights into how ecosystem and plant traits influence endophytic fungi, and how these microorganisms, in turn, affect host and ecosystem functions. It is imperative to continue the efforts to understand the degree to which apparent patterns of host colonization and selection for diverse endophytes are dictated by the host, the endophyte or other ecological mechanisms.

## ACKNOWLEDGEMENTS

We thank botanist Pedro Juarez-Retana for the plant identification, Efraín Escudero-Leyva for his help in sample collection, and Wilber Alvarado for assistance in the field. Dr. Gloriana Chaverri and Dr. Óscar Quirós from Fundación ProSur, for the access to sites in Golfito and lab space during fieldtrips. Samples were collected under the Institutional Biodiversity Commission (University of Costa Rica) resolutions VI07723-2018 and VI5152-2016, and Costa Rica’s National System of Conservation Areas (SINAC), permit N° ACG-PI-017-2017. Thanks to Dr. Maile Neel for her feedback; and Dr. David Zelený and Joseph Murphy for help with bioinformatics.

## SUPPLEMENTARY INFORMATION

Supplementary information is publicly available in Zenodo https://doi.org/10.5281/zenodo.13892068

**Supplementary Table S1.** List of putative plant species collected based on morphology. In bold are the species collected twice.

Supplementary Methods S2.

**Supplementary Table S3**. Taxonomic assignment (7 ranks) of all ASVs identified in the dataset, plus their functional traits, can be found publicly available on the Github repository https://github.com/humb15/RuFEnTroFo.

**Supplementary Figure S4**. Species accumulation and diversity curves for endophytes sampled from (A) young leaves (YL), mature leaves (ML) and sapwood (Sa) tissues, and; (B) Golfito and Guanacaste forests. Metrics include richness (q = 0), Shannon (q = 1) and Simpson (q = 2).

**Supplementary Figure S5**. NMDS of Bray Curtis distances showing the distribution of endophytic community assemblages according to (A) genus within tribes and (B) tribes within subfamilies.

**Supplementary Table S6**. List of identified fungal families among endophyte communities.

**Supplementary Figure S7**. CCA analysis for correlation between Amplicon Single Variants distribution and environmental variables: Elevation, Annual Mean Temperature and Annual Precipitation). Red crosses represent ASVs and black circles represent samples. p-values: Elevation 0.001, AMT 0.073, AP 0.119; CCA1 0.007, CCA2 0.143.

**Supplementary Table S8**. Mantel correlation values for community dissimilarity and geographic and environmental factors. Based on Spearman, p-value reported.

**Supplementary Figure S9**. Distance decay of taxonomic similarity along spatial distance, every point represents pairwise similarities between samples. Panels A: combined; B: Golfito; C: Guanacaste.

**Supplementary Figure S10**. Main lifestyles (primary and secondary) of the genera identified among the top 12 families.

## DATA AVAILABILITY

The resulting raw data (fastq files) and supporting metadata are deposited and publicly available in GenBank under the BioProject identifier PRJNA889378.

## AUTHOR CONTRIBUTIONS

Conceptualization: P.C., H.C., J.S.; Formal analysis: H.C.; Funding acquisition: P.C., J.S.; Investigation: H.C.; Methodology: H.C., P.C., S.Y.; Project administration: P.C., S.Y.; Resources: P.C.; Supervision: P.C., S.Y.; Validation: P.C., S.Y., J.S., H.C.; Visualization: H.C., S.Y; Writing: H.C., J.S., S.Y., <article>P</article>.C.

