## Supplementary Information for "Exploring Rubiaceae fungal endophytes across contrasting tropical forests, tree tissues, and developmental stages"

**Supplementary Table S1.** List of putative plant species collected based on morphology. In **bold** are the species collected twice.

| Golfito | Guanacaste |
|---|---|
| *Chimarrhis latifolia* | ***Alibertia edulis*** |
| *Chomelia microloba* | *Allenanthus erythrocarpus* |
| *Coussarea loftonii* | *Arachnothryx costaricensis* |
| *Coussarea talamancana* | *Bertiera bracteosa* |
| ***Duroia costaricensis*** | *Chomelia costaricensis* |
| *Faramea tamberlikiana* | *Cosmibuena macrocarpa* |
| *Gonzalagunia osaensis* | *Genipa americana* |
| *Isertia haenkeana* | *Guettarda macrosperma* |
| *Macrocnemum roseum* | *Palicourea adusta* |
| *Notopleura polyphlebia* | *Palicourea eurycarpa* |
| *Posoqueria grandiflora* | *Pentagonia costaricensis* |
| *Posoqueria latifolia* | *Psychotria berteroana* |
| *Psychotria brachiata* | *Psychotria callophyla* |
| *Psychotria chagrensis* | *Psychotria guianensis* |
| *Psychotria mortoniana* | *Psychotria horizontalis* |
| ***Psychotria panamensis*** | *Psychotria marginata* |
| *Psychotria solitodinum* | *Psychotria pubescens* |
| *Ronabea emética* | *Psychotria subssesilis* |
| *Ronabea latifolia* | *Psychotria valerioana* |
| *Rudgea cornifolia* | *Randia aculeata* |
| *Rustia costaricensis* | *Randia grandifolia* |
| *Simira maxonii* | *Randia lasiantha* |
| *Sommera donnell-smithii* | |
| *Tocoyena pittieri* | |
| *Warszewiczia coccinea* | |

**Supplementary Methods S2**

Rarefaction curves were calculated using the 'iNEXT' function from the iNEXT package v 3.0.0 (Hsieh et al., 2016, 2022) and species accumulation curves were plotted using the 'ggiNEXT' function, to determine the if the sampling effort was enough to accurately assess the fungal endophytic biodiversity present in the samples.

Mantel correlation coefficients (using the 'mantel' function in the vegan package, "spearman" method and 1000 permutations) were used to assess the relationship between environmental variables and endophyte abundance, and to explore endophyte dispersal limitation, as a measure of the strength of distance-decay relationships. Distance matrices for environmental variables were generated using the 'dist' function (method = "euclidean") and for geographic distance the 'distm' (Haversine method) from the 'geosphere' package, v 1.5.18 (Hijmans et al., 2022). Endophyte abundance matrix based on Bray-Curtis dissimilarity index, was calculated with the 'vegdist' function. We also performed an additional analysis using a distance matrix of scaled data that included all environmental variables. To assess the significance of the results, we used the 'envfit' function.

Constrained correspondence analysis (a.k.a. canonical correspondence analysis) was conducted to explore the relationship between species abundances and three environmental variables: elevation, AMP, and AT. The analysis was performed using the vegan package in R, with the 'cca' function to generate the CCA model. To select the most significant environmental variables, we used the 'ordistep' function and 'anova.cca' to test for the significance of the selected variables in the final model. This approach allowed us to assess the extent to which environmental variables explain the variation in species abundances and identify the most important environmental predictors of community structure.

To characterize the functional traits of the endophytes identified in our study, we used the FungalTraits database (Põlme et al., 2020). Specifically, we made the assignment based on the primary and secondary lifestyles.

**Supplementary Table S3.** Taxonomic assignment (7 ranks) of all ASVs identified in the dataset, plus their functional traits, can be found publicly available on the Github repository https://github.com/humb15/RuFEnTroFo.

**Supplementary Figure S4.** Species accumulation and diversity curves for endophytes sampled from **(A)** young leaves (YL), mature leaves (ML) and sapwood (Sa) tissues, and; **(B)** Golfito and Guanacaste forests. Metrics include richness (q = 0), Shannon (q = 1) and Simpson (q = 2).

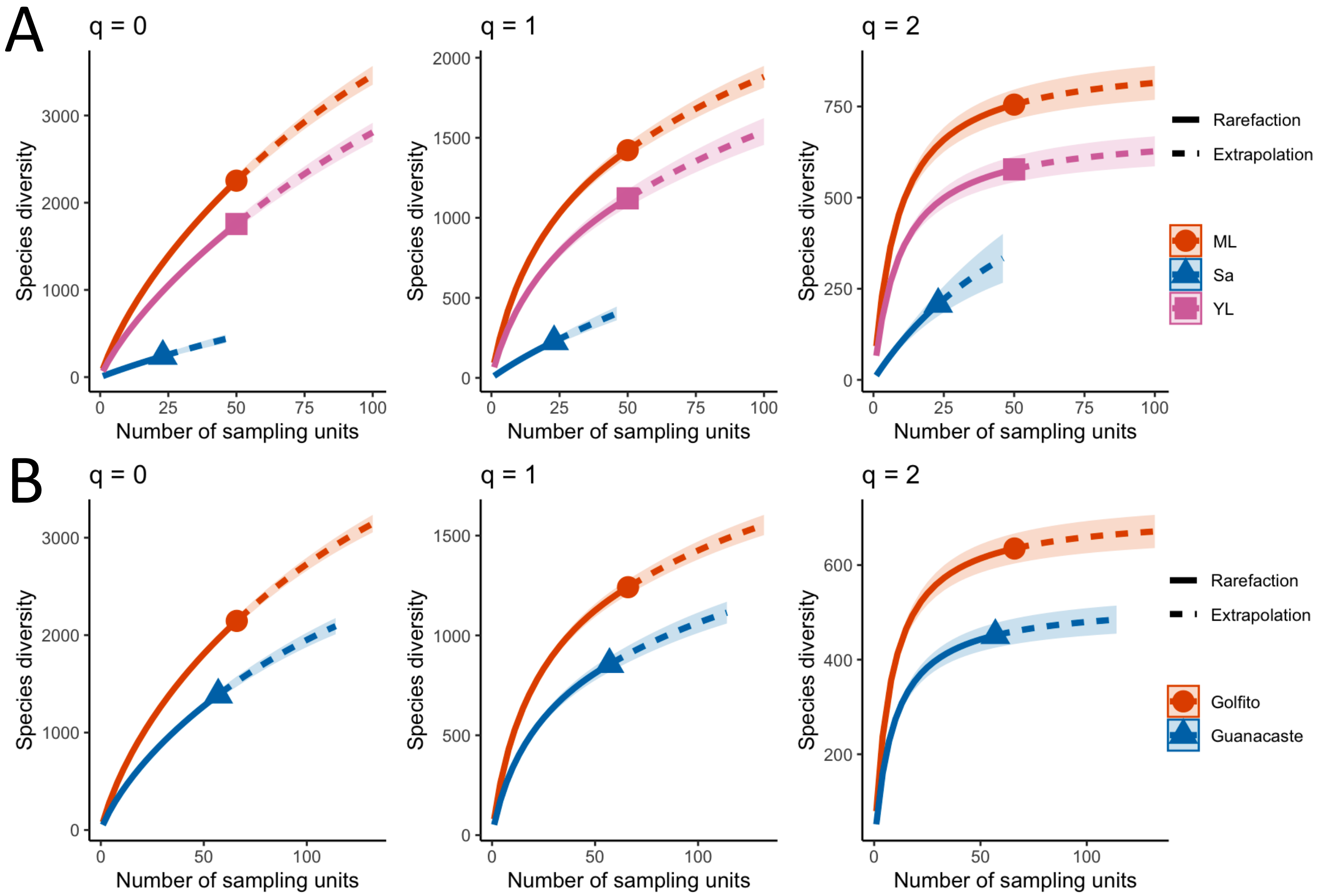

**Supplementary Figure S5.** NMDS of Bray Curtis distances showing the distribution of endophytic community assemblages according to (A) genus within tribes and (B) tribes within subfamilies.

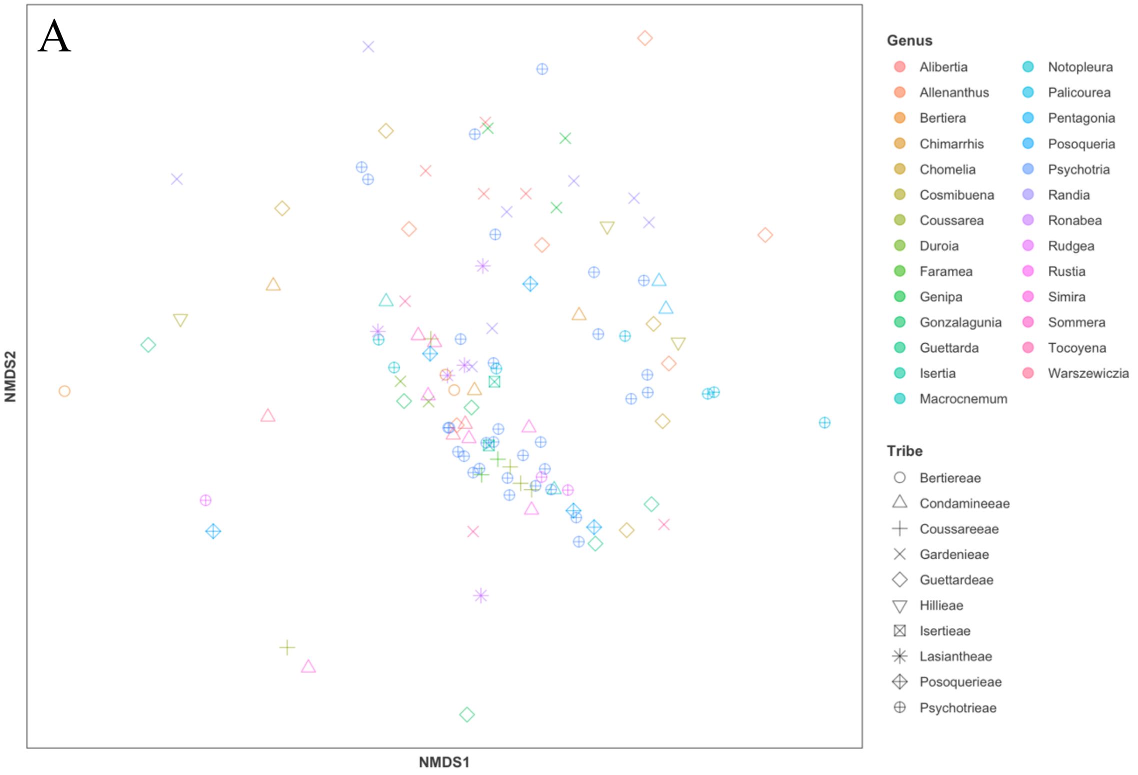


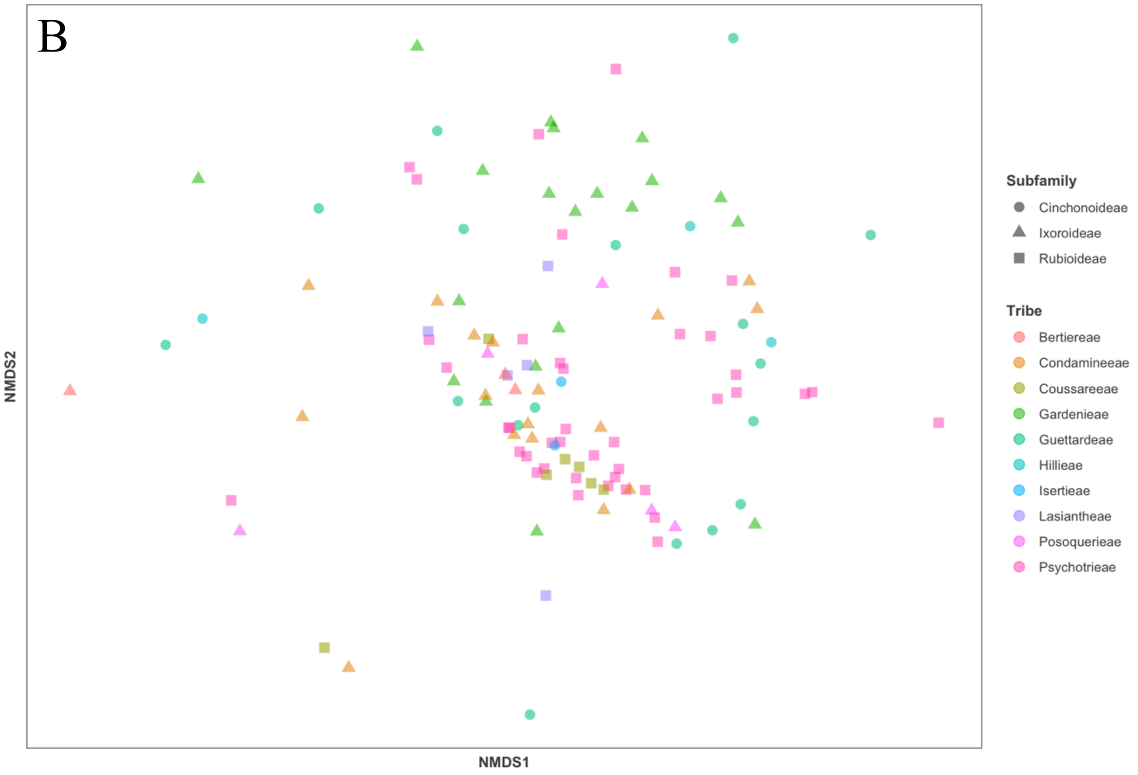

**Supplementary Table S6.** List of identified fungal families among endophyte communities.

| | Family |
|---|---|
| **1** | *Agaricaceae* |
| **2** | *Agaricomycetes_fam_Incertae_sedis* |
| **3** | *Amphisphaeriaceae* |
| **4** | *Amplistromataceae* |
| **5** | *Anungitiomycetaceae* |
| **6** | *Apiosporaceae* |
| **7** | *Aspergillaceae* |
| **8** | *Aureobasidiaceae* |
| **9** | *Auriculariaceae* |
| **10** | *Australiascaceae* |
| **11** | *Beltraniaceae* |
| **12** | *Biatriosporaceae* |
| **13** | *Bionectriaceae* |
| **14** | *Bolbitiaceae* |
| **15** | *Boliniales_fam_Incertae_sedis* |
| **16** | *Botryobasidiaceae* |
| **17** | *Botryosphaeriaceae* |
| **18** | *Brachybasidiaceae* |
| **19** | *Bulleribasidiaceae* |
| **20** | *Cenangiaceae* |
| **21** | *Ceratobasidiaceae* |
| **22** | *Chaetosphaeriaceae* |
| **23** | *Chaetothyriales_fam_Incertae_sedis* |
| **24** | *Chrysozymaceae* |
| **25** | *Cladosporiaceae* |
| **26** | *Clathraceae* |
| **27** | *Clavicipitaceae* |
| **28** | *Coniocessiaceae* |
| **29** | *Coniophoraceae* |
| **30** | *Cordycipitaceae* |
| **31** | *Corynesporascaceae* |
| **32** | *Cucurbitariaceae* |
| **33** | *Cuniculitremaceae* |
| **34** | *Cyphellophoraceae* |
| **35** | *Cystobasidiaceae* |
| **36** | *Cystofilobasidiaceae* |
| **37** | *Debaryomycetaceae* |
| **38** | *Dermateaceae* |

| | Family |
|---|---|
| **79** | *Malasseziaceae* |
| **80** | *Massarinaceae* |
| **81** | *Melanconidaceae* |
| **82** | *Meripilaceae* |
| **83** | *Meruliaceae* |
| **84** | *Microbotryomycetes_fam_Incertae_sedis* |
| **85** | *Microdochiaceae* |
| **86** | *Microstromatales_fam_Incertae_sedis* |
| **87** | *Morosphaeriaceae* |
| **88** | *Mycosphaerellaceae* |
| **89** | *Myriangiaceae* |
| **90** | *Nectriaceae* |
| **91** | *Neodevriesiaceae* |
| **92** | *Neomassarinaceae* |
| **93** | *Niessliaceae* |
| **94** | *Ophiocordycipitaceae* |
| **95** | *Ophiostomataceae* |
| **96** | *Peniophoraceae* |
| **97** | *Pezizomycotina_fam_Incertae_sedis* |
| **98** | *Phaeococcomycetaceae* |
| **99** | *Phaeosphaeriaceae* |
| **100** | *Phaeotremellaceae* |
| **101** | *Phanerochaetaceae* |
| **102** | *Phyllostictaceae* |
| **103** | *Physciaceae* |
| **104** | *Plectosphaerellaceae* |
| **105** | *Pleosporaceae* |
| **106** | *Pleosporales_fam_Incertae_sedis* |
| **107** | *Pluteaceae* |
| **108** | *Polyporaceae* |
| **109** | *Psathyrellaceae* |
| **110** | *Pseudoberkleasmiaceae* |
| **111** | *Pterulaceae* |
| **112** | *Pyrenulaceae* |
| **113** | *Ramalinaceae* |
| **114** | *Rhynchogastremataceae* |
| **115** | *Rhytismataceae* |
| **116** | *Rickenellaceae* |

| | |
|---|---|
| **39** | *Diaporthaceae* |
| **40** | *Dictyosporiaceae* |
| **41** | *Didymellaceae* |
| **42** | *Didymosphaeriaceae* |
| **43** | *Dissoconiaceae* |
| **44** | *Dothioraceae* |
| **45** | *Elsinoaceae* |
| **46** | *Entolomataceae* |
| **47** | *Erythrobasidiaceae* |
| **48** | *Exidiaceae* |
| **49** | *Fasciatisporaceae* |
| **50** | *Filobasidiaceae* |
| **51** | *Fomitopsidaceae* |
| **52** | *Ganodermataceae* |
| **53** | *Geastraceae* |
| **54** | *Glomerellaceae* |
| **55** | *Glomerellales_fam_Incertae_sedis* |
| **56** | *Halosphaeriaceae* |
| **57** | *Helminthosphaeriaceae* |
| **58** | *Helotiaceae* |
| **59** | *Herpotrichiellaceae* |
| **60** | *Hyaloriaceae* |
| **61** | *Hyaloscyphaceae* |
| **62** | *Hydnodontaceae* |
| **63** | *Hymenochaetaceae* |
| **64** | *Hymenochaetales_fam_Incertae_sedis* |
| **65** | *Hyphodermataceae* |
| **66** | *Hypocreaceae* |
| **67** | *Hypocreales_fam_Incertae_sedis* |
| **68** | *Hyponectriaceae* |
| **69** | *Inocybaceae* |
| **70** | *Irpicaceae* |
| **71** | *Lachnocladiaceae* |
| **72** | *Lasiosphaeriaceae* |
| **73** | *Lecanoromycetes_fam_Incertae_sedis* |
| **74** | *Lentitheciaceae* |
| **75** | *Lophiostomataceae* |
| **76** | *Lophiotremataceae* |
| **77** | *Lycoperdaceae* |
| **78** | *Magnaporthaceae* |

| | |
|---|---|
| **117** | *Saccharomycetales_fam_Incertae_sedis* |
| **118** | *Sarcoscyphaceae* |
| **119** | *Schizoparmaceae* |
| **120** | *Schizophyllaceae* |
| **121** | *Schizoporaceae* |
| **122** | *Sclerotiniaceae* |
| **123** | *Septobasidiaceae* |
| **124** | *Sordariaceae* |
| **125** | *Sordariomycetes_fam_Incertae_sedis* |
| **126** | *Sporidiobolaceae* |
| **127** | *Sporocadaceae* |
| **128** | *Sporormiaceae* |
| **129** | *Stachybotryaceae* |
| **130** | *Steccherinaceae* |
| **131** | *Stereaceae* |
| **132** | *Stictidaceae* |
| **133** | *Strophariaceae* |
| **134** | *Sulcatisporaceae* |
| **135** | *Sympoventuriaceae* |
| **136** | *Teloschistaceae* |
| **137** | *Teratosphaeriaceae* |
| **138** | *Tetraplosphaeriaceae* |
| **139** | *Thyridariaceae* |
| **140** | *Tilachlidiaceae* |
| **141** | *Togniniaceae* |
| **142** | *Tremellaceae* |
| **143** | *Tremellales_fam_Incertae_sedis* |
| **144** | *Trichocomaceae* |
| **145** | *Tricholomataceae* |
| **146** | *Trichosphaeriaceae* |
| **147** | *Trimorphomycetaceae* |
| **148** | *Tulasnellaceae* |
| **149** | *Tylosporaceae* |
| **150** | *Valsaceae* |
| **151** | *Vuilleminiaceae* |
| **152** | *Wallemiaceae* |
| **153** | *Wiesneriomycetaceae* |
| **154** | *Wrightoporiaceae* |
| **155** | *Xylariaceae* |
| **156** | *Xylariales_fam_Incertae_sedis* |

**Supplementary Figure S7.** CCA analysis for correlation between Amplicon Single Variants distribution and environmental variables: Elevation, Annual Mean Temperature and Annual Precipitation). Red crosses represent ASVs and black circles represent samples. p-values: Elevation 0.001, AMT 0.073, AP 0.119; CCA1 0.007, CCA2 0.143.

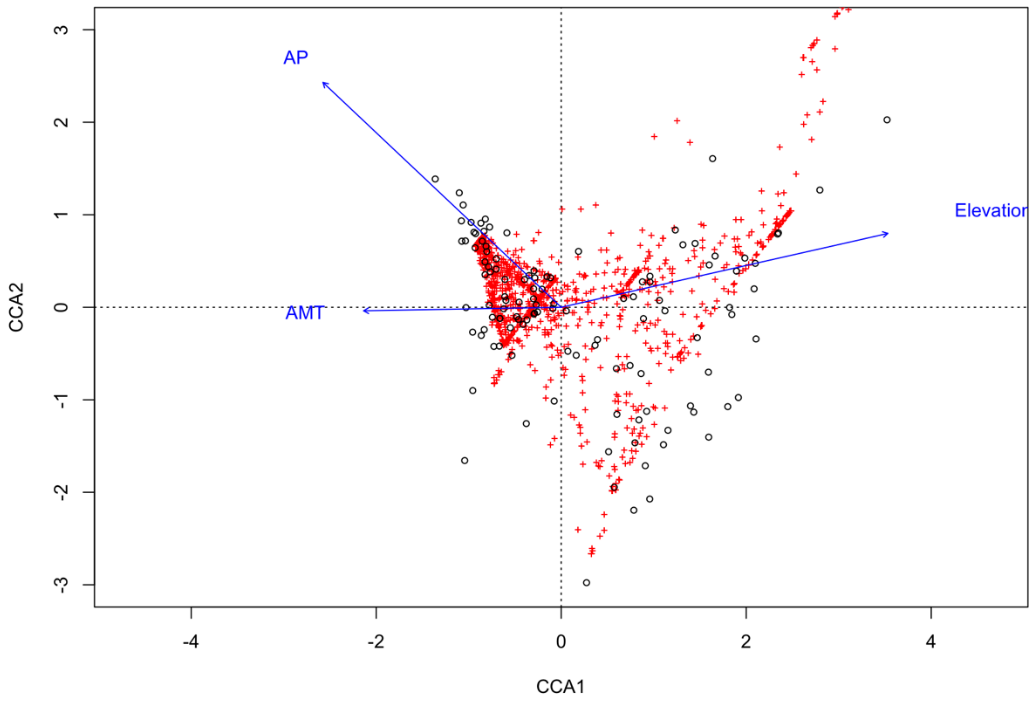


**Supplementary Table S8.** Mantel correlation values for community dissimilarity and geographic and environmental factors. Based on Spearman, p-value reported.

| Factor | | Mantel statistic R | p-value (Significance) |
|---|---|---|---|
| Distance (combined) | | 0.2434 | 0.000999 |
| | *Golfito* | 0.1875 | 0.002997 |
| | *Guanacaste* | 0.0681 | 0.17682 |
| Environment (combined) | | 0.1496 | 0.001998 |
| | *Temperature* | 0.1258 | 0.001998 |
| | *Elevation* | 0.1998 | 0.000999 |
| | *Precipitation* | 0.1889 | 0.000999 |

**Supplementary Figure S9.** Distance decay of taxonomic similarity along spatial distance, every point represents pairwise similarities between samples. Panels A: combined; B: Golfito; C: Guanacaste.

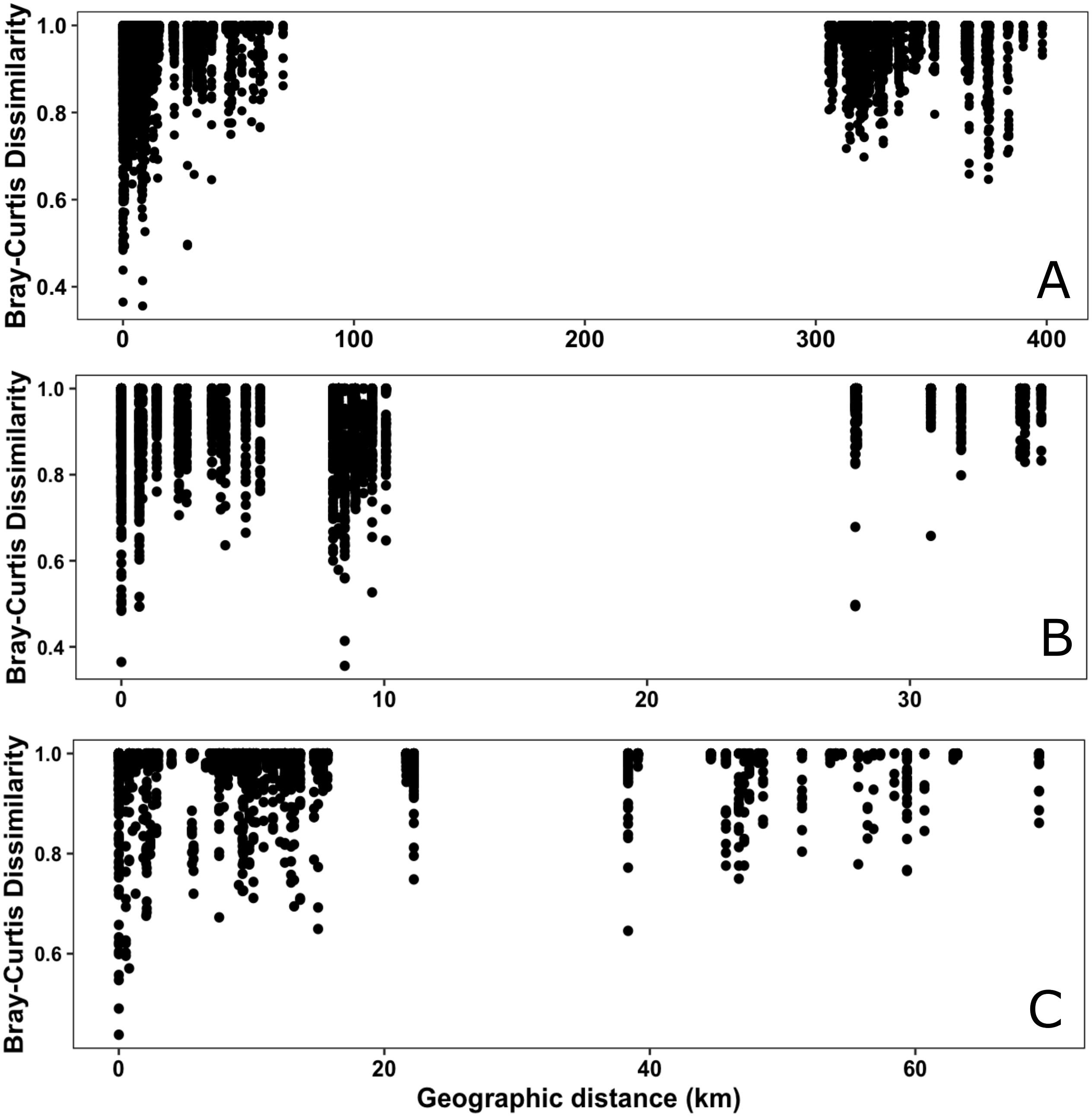

**Supplementary Figure S10.** Main lifestyles (primary and secondary) of the genera identified among the top 12 families.

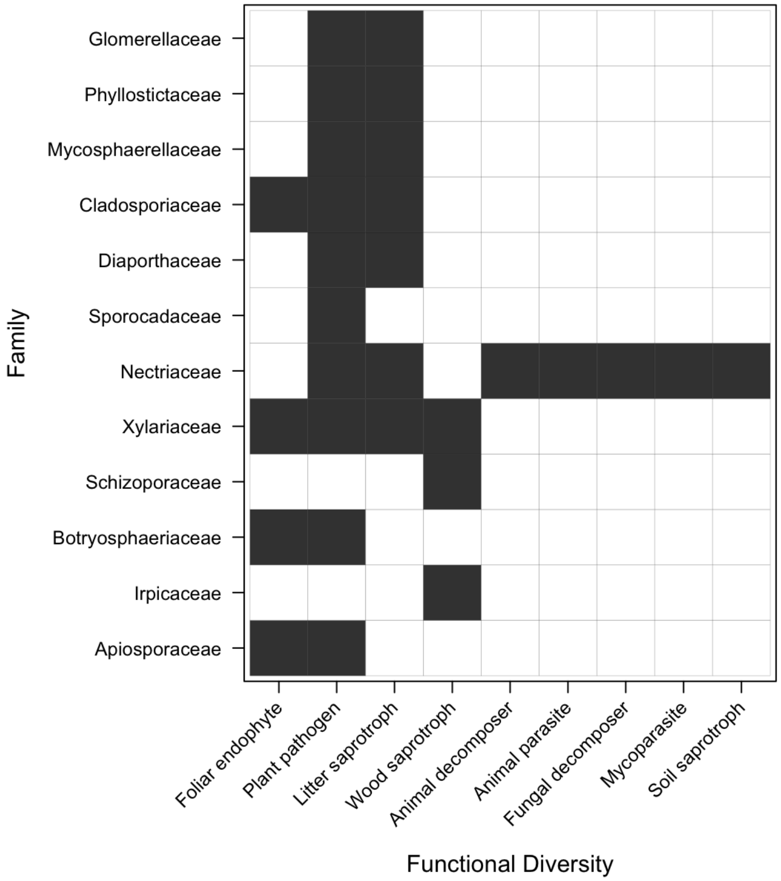
